# Supplier-associated variation in motor performance and cerebrospinal fluid-contacting neuron anatomy in C57BL/6 mice

**DOI:** 10.64898/2026.11.24.769017

**Authors:** Meili J. Luther, Kathryn E. Stanchak, David J. Perkel

## Abstract

Cerebrospinal fluid-contacting neurons (CSF-cNs) are an evolutionarily conserved population of spinal sensory neurons positioned around the central canal and characterized by expression of the ion channel PKD2L1. CSF-cNs contribute to locomotor control, but little is known about variation in this sensory system among commonly used laboratory mouse populations. We compared motor performance and CSF-cN organization in closely related C57BL/6 mice obtained from The Jackson Laboratory (Jackson), Charles River Laboratories (Charles River), and Taconic Biosciences (Taconic). Motor performance differed among suppliers, with Charles River mice performing better than Taconic mice on balance beam and rotarod tasks, whereas grid walking performance did not differ significantly among groups. CSF-cN organization also differed significantly among suppliers. Jackson mice had the greatest abundance of PKD2L1+ cells and canal-contacting cells, and a greater proportion of PKD2L1+ cells contacted the central canal in Jackson mice than in Charles River or Taconic mice. CSF-cNs were also more abundant in cervical than lumbar spinal cord regions, whereas the proportion contacting the canal showed only a trend toward a regional difference. Individual CSF-cN measures did not show significant associations with motor performance. These findings demonstrate significant supplier-associated variation in both motor performance and the organization of the CSF-cN anatomy and show that greater CSF-cN abundance does not necessarily correspond to better motor performance.

**SIGNIFICANCE STATEMENT:** Closely related C57BL/6 mouse populations obtained from different commercial suppliers can differ in motor performance, but whether their spinal sensory anatomy also varies among these populations is less understood. We show that supplier-associated differences in motor performance occur alongside substantial differences in the abundance and central canal organization of PKD2L1+ cerebrospinal fluid-contacting neurons (CSF-cNs). CSF-cN abundance also differs between cervical and lumbar spinal cord. However, greater CSF-cN abundance does not directly correspond to better motor performance, suggesting that cell number alone does not determine the functional contribution of this sensory population. These findings identify commercial source and spinal region as important sources of variation in the mouse CSF-cN system.

## INTRODUCTION

The spinal cord contains a distinct subpopulation of sensory neurons that directly interact with the cerebrospinal fluid (CSF), called cerebrospinal fluid-contacting neurons (CSF-cNs). CSF-cNs were first described by Agduhr and Kolmer in the early twentieth century and have since been identified across a wide range of vertebrate species (Agduhr, 1922; Kolmer, 1931; Djenoune et al., 2014; Djenoune and Wyart, 2017; Wyart et al., 2023). CSF-cNs are positioned within or immediately adjacent to the ependymal layer surrounding the central canal. They extend an apical process toward the canal that terminates in a bulbous protrusion into the canal containing cilia and microvilli, and therefore in direct contact with the CSF (Orts-Del’Immagine et al., 2012; Djenoune et al., 2014; Djenoune and Wyart, 2017; Wyart et al., 2023; Crozat et al., 2025; Vitar et al., 2026). A defining molecular feature of CSF-cNs is the expression of polycystic kidney disease 2-like-1 (PKD2L1), a member of the transient receptor potential (TRP) ion channel family (Delmas, 2004). PKD2L1 is enriched in the apical compartment of CSF-cNs and has been established as a selective marker of CSF-cNs in zebrafish, mice, and primates (Orts-Del’Immagine et al., 2012; Djenoune et al., 2014; Djenoune and Wyart, 2017; Wyart et al., 2023).

Much of our understanding of CSF-cN mechanosensation function comes from zebrafish (Fidelin et al., 2015; Böhm et al., 2016; Hubbard et al., 2016; Djenoune and Wyart, 2017). CSF-cNs respond to bending of the spinal cord and relay sensory information to locomotor circuits (Fidelin et al., 2015; Böhm et al., 2016; Hubbard et al., 2016; Djenoune and Wyart, 2017). This mechanosensation depends on PKD2L1, as loss of pkd2l1 disrupts CSF-cN responses to spinal bending and reduces tail-beat frequency during swimming (Böhm et al., 2016). CSF-cNs form GABAergic synapses with premotor interneurons and motor neurons, providing a pathway through which sensory information can influence spinal locomotor circuits (Fidelin et al., 2015; Böhm et al., 2016). Silencing CSF-cNs disrupts postural control during locomotion (Hubbard et al., 2016), and rostral CSF-cNs project to the hindbrain and contribute to locomotor speed and postural stabilization (Wu et al., 2021). CSF-cNs also detect chemical properties of their environment. In lampreys, CSF-cNs respond to changes in extracellular pH through acid-sensing ion channels (Jalalvand et al., 2016). Mouse spinal CSF-cNs are likewise sensitive to extracellular pH through PKD2L1 and acid-sensing ion channels (Crozat et al., 2025). PKD2L1 channels localized to the apical compartment of mouse CSF-cNs are particularly sensitive to extracellular pH, further supporting a chemosensory role for these neurons (Vitar et al., 2026). These studies establish CSF-cNs as multimodal sensory neurons that detect mechanical and chemical changes in their environment and influence motor output across vertebrates.

Studies in mice have directly linked CSF-cNs to locomotor function. CSF-cNs extend long rostrocaudal axons within the spinal cord and form synaptic connections with premotor interneurons, motor neurons, and other CSF-cNs (Orts-Del’Immagine et al., 2012; Gerstmann et al., 2022; Nakamura et al., 2023). Ablation of Pkd2l1-expressing CSF-cNs increases foot-placement errors during skilled locomotor tasks while leaving general locomotor activity and basic gait largely intact (Gerstmann et al., 2022). Chemogenetic inhibition of Pkd2l1-expressing CSF-cNs reduces maximal treadmill speed and step frequency and increases stride length without disrupting left-right limb coordination (Nakamura et al., 2023). Thus, CSF-cNs contribute to specific aspects of mammalian locomotor control.

Motor performance can differ among closely related C57BL/6 populations obtained from different commercial suppliers. Direct comparisons of C57BL/6J, C57BL/6NCrl, and C57BL/6NTac mice have identified differences in motor coordination and other behavioral phenotypes (Bryant et al., 2008; Mekada et al., 2009; Mekada and Yoshiki, 2021). CSF-cN organization may also vary among C57BL/6 populations. Tonelli Gombalová et al. (2020) reported marked differences in CSF-cN positioning between C57BL/6J and C57BL/6N mice: most PKD2L1+ CSF-cNs examined in C57BL/6N mice occupied an ectopic ventral position, whereas ectopic cells represented only a small population in C57BL/6J mice. We hypothesized that C57BL/6 mice obtained from different commercial suppliers would exhibit differences in motor performance accompanied by variation in the abundance and organization of spinal CSF-cNs. To test this hypothesis, we compared mice obtained from The Jackson Laboratory (Jackson), Charles River Laboratories (Charles River), and Taconic Biosciences (Taconic).

We assessed motor performance and quantified the abundance and central canal organization of PKD2L1+ CSF-cNs in cervical and lumbar spinal cord regions from the same animals. We also examined whether anatomical measurements differed between spinal regions and whether individual variation in CSF-cN anatomy was associated with motor performance. Together, these analyses allowed us to characterize supplier-associated differences in motor behavior and spinal CSF-cN organization and evaluate potential relationships between these phenotypes.

## RESULTS

### Motor performance differs among commercial suppliers

We first asked whether motor performance differed among mice obtained from the three suppliers. We assessed performance using balance beam, grid walking, and rotarod tasks (Fig. 1).

**Figure 1.**
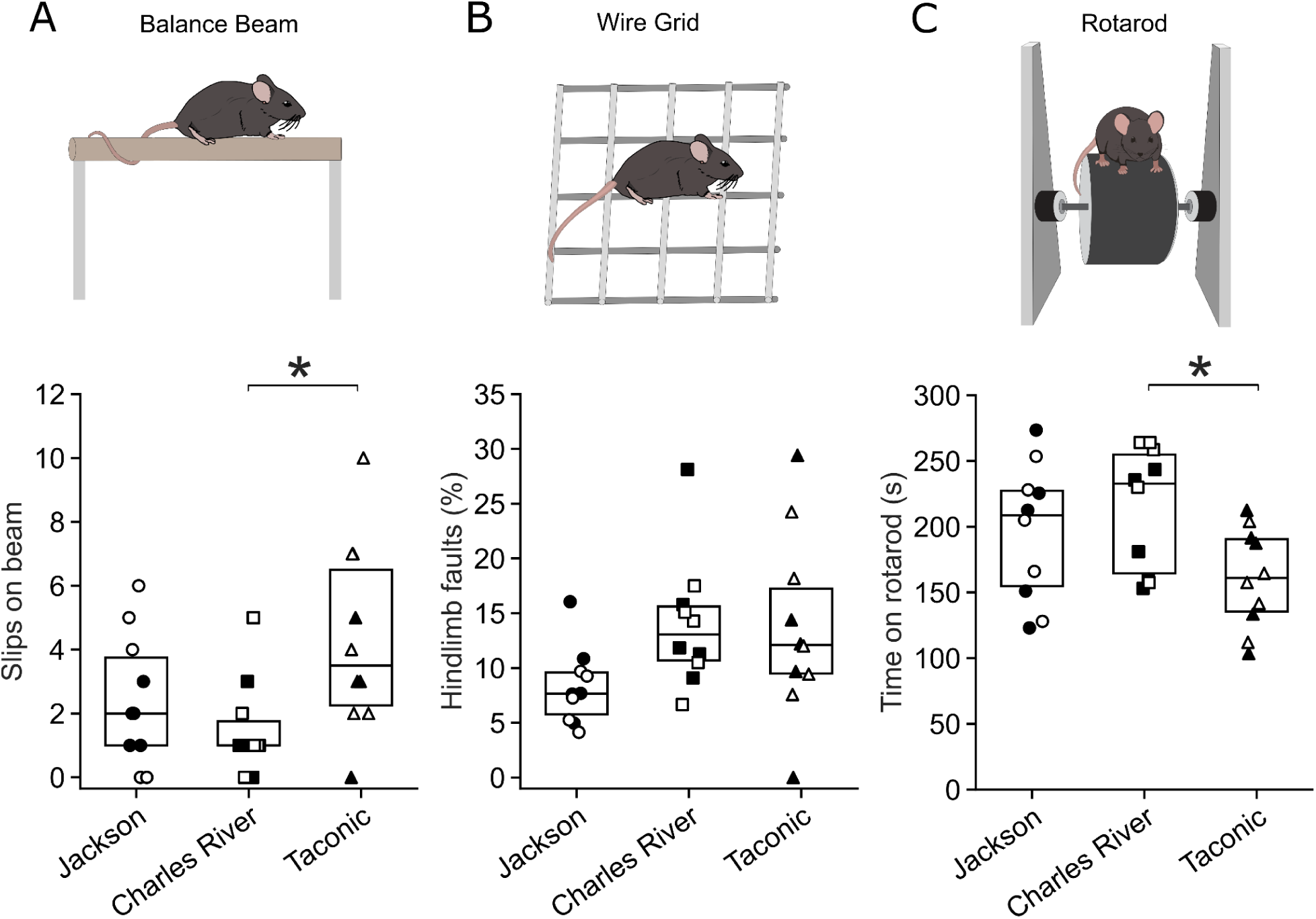
Motor performance differs among commercial suppliers. Motor performance was assessed in mice obtained from The Jackson Laboratory, Charles River Laboratories, and Taconic Biosciences using balance beam, grid walk, and rotarod tasks. Schematics of behavioral tests (top row). (A) Number of slips on the balance beam. Jackson: Q1 = 1.00, Q2 = 2.00, Q3 = 3.75; Charles River: Q1 = 1.00, Q2 = 1.00, Q3 = 1.75; Taconic: Q1 = 2.25, Q2 = 3.50, Q3 = 6.50. (B) Percentage of hindlimb faults on the grid. Jackson: Q1 = 5.76%, Q2 = 7.66%, Q3 = 9.59%; Charles River: Q1 = 10.69%, Q2 = 13.06%, Q3 = 15.62%; Taconic: Q1 = 9.49%, Q2 = 12.10%, Q3 = 17.23%. (C) Time spent on the rotarod. Jackson: Q1 = 154.75 s, Q2 = 208.75 s, Q3 = 227.38 s; Charles River: Q1 = 164.50 s, Q2 = 232.75 s, Q3 = 254.75 s; Taconic: Q1 = 135.50 s, Q2 = 161.00 s, Q3 = 190.50 s. Box plots show the median (Q2), first quartile (Q1), and third quartile (Q3), with individual animals shown as overlaid symbols. Jackson mice are represented by circles, Charles River mice by squares, and Taconic mice by triangles. Male mice are shown with filled symbols and female mice with open symbols. Asterisks indicate significant pairwise differences following Tukey’s HSD test: *p < 0.05.

The number of foot slips on the balance beam differed significantly among mice from the different suppliers (Fig. 1A; one-way ANOVA, *F*(2,27) = 3.97, *p* = 0.0308). Charles River mice made the fewest slips on average (1.5 ± 1.5), followed by Jackson mice (2.4 ± 2.07), whereas Taconic mice made the most slips (4.3 ± 2.98). Post hoc Tukey tests identified a significant difference between Charles River and Taconic mice (*p* = 0.0269), while Jackson mice did not differ significantly from either other group.

Grid performance did not differ significantly among suppliers (Fig. 3B; *F*(2,27) = 2.63, *p* = 0.0908). Jackson mice had the lowest percentage of hindlimb faults (8.28 ± 3.48%), compared with Charles River (14.02 ± 5.95%) and Taconic mice (13.71 ± 8.45%), but no pairwise comparison reached statistical significance.

Rotarod performance also differed significantly among suppliers (Fig. 3C; *F*(2,27) = 3.55, *p* = 0.0427). Charles River mice remained on the rotarod longest on average (214s ± 46.66 s), followed by Jackson mice (196.6 ± 52.11 s) and Taconic mice (160.8 ± 38.01 s). Charles River mice performed significantly better than Taconic mice (*p* = 0.0370, Tukey’s HSD), whereas the remaining pairwise comparisons were not significant. These results indicate that motor performance varies among commercial suppliers, although the pattern of differences depends on the behavioral task.

Because CSF-cNs contribute to locomotor control and previous work identified differences in CSF-cN organization among C57BL/6 populations, we next asked whether CSF-cN organization also differed among these commercially sourced mice.

### CSF-cN abundance and central canal organization differ among commercial suppliers

We next measured four CSF-cN anatomical variables in cervical and lumbar spinal cord sections from male and female mice obtained from the three suppliers (Fig. 2D-G). To evaluate the effects of supplier, spinal region, and sex while accounting for paired regional measurements within animals, we performed mixed-design repeated-measures ANOVAs, with supplier and sex as between-subject factors and spinal region as a within-subject factor.

**Figure 2.**
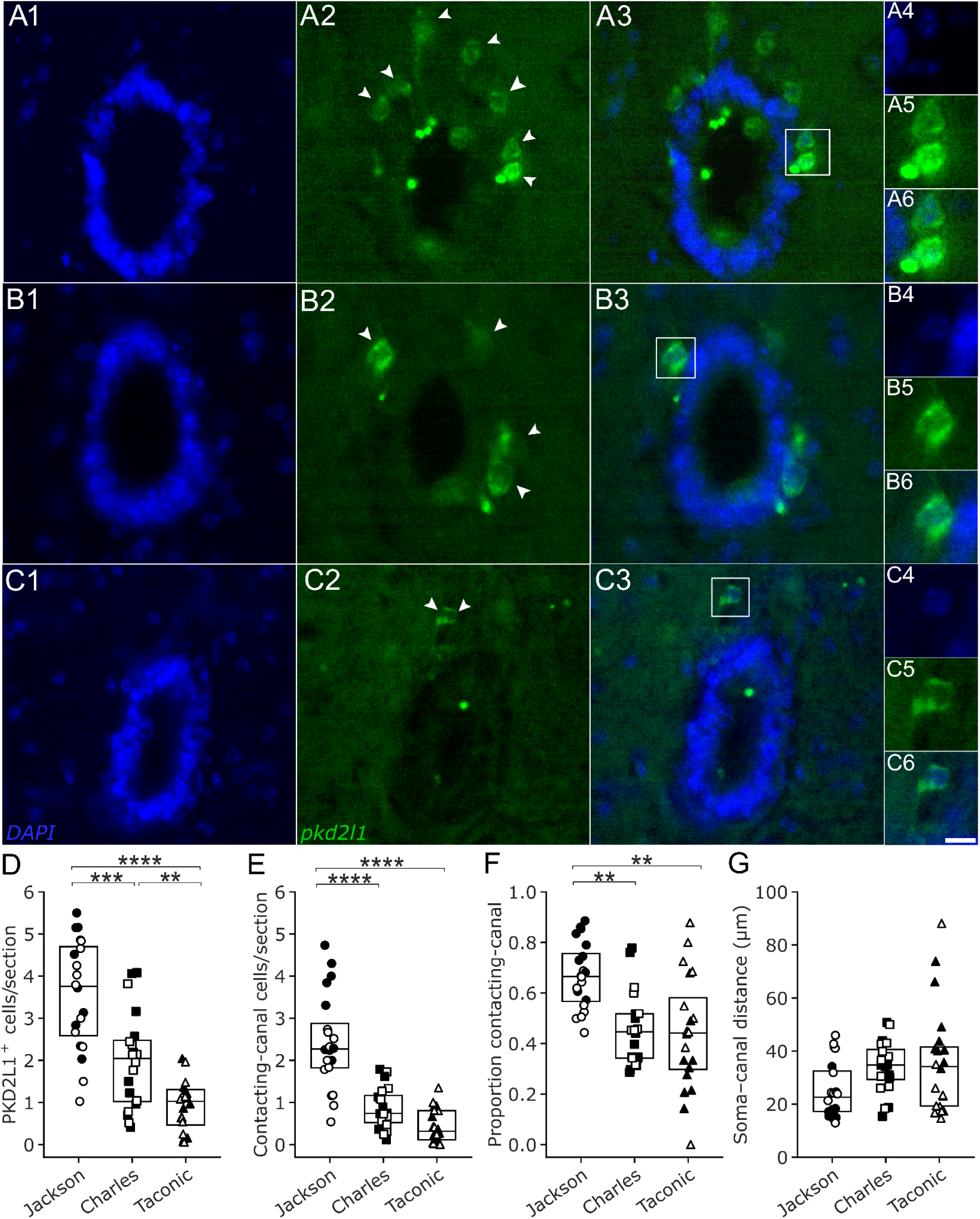
PKD2L1+ CSF-cN expression and distribution across commercial suppliers. Representative spinal cord sections from Jackson (A1-A6), Charles River (B1-B6), and Taconic (C1-C6) mice immunolabeled for PKD2L1+ expressing cells. A1, B1, C1, DAPI channel; A2, B2, C2, PKD2L1 channel; A3, B3, C3, merged images. A4, B4, C4, higher-magnification DAPI images of the central canal region; A5, B5, C5, higher-magnification PKD2L1 images; A6, B6, C6, higher-magnification merged images. The scale bar is 10µm. (D) Average number of PKD2L1+ cells per section. Jackson: Q2 = 3.76, Q1 = 2.59, Q3 = 4.70; Charles River: Q2 = 2.04, Q1 = 1.02, Q3 = 2.48; Taconic: Q2 = 1.03, Q1 = 0.47, Q3 = 1.31. (E) Average number of PKD2L1+ cells per section contacting the central canal. Jackson: Q2 = 2.27, Q1 = 1.82, Q3 = 2.88; Charles River: Q2 = 0.74, Q1 = 0.52, Q3 = 1.17; Taconic: Q2 = 0.32, Q1 = 0.11, Q3 = 0.81. (F) Average distance of PKD2L1+ cell somata from the central canal. Jackson: Q2 = 22.60 µm, Q1 = 17.21 µm, Q3 = 32.49 µm; Charles River: Q2 = 34.76 µm, Q1 = 29.28 µm, Q3 = 40.61 µm; Taconic: Q2 = 34.16 µm, Q1 = 19.31 µm, Q3 = 41.53 µm. (G) Proportion of PKD2L1+ cells contacting the central canal. Jackson: Q2 = 0.67, Q1 = 0.57, Q3 = 0.76; Charles River: Q2 = 0.45, Q1 = 0.34, Q3 = 0.52; Taconic: Q2 = 0.44, Q1 = 0.30, Q3 = 0.58. Box plots show the median (Q2), first quartile (Q1), and third quartile (Q3). Jackson mice are represented by circles, Charles River mice by squares, and Taconic mice by triangles (n = 30 mice). Lumbar measurements are shown with filled symbols and cervical measurements with open symbols. Asterisks indicate significant pairwise differences between suppliers following Tukey’s HSD test: *p < 0.05, **p < 0.01, ***p < 0.001, and ****p < 0.0001.

The number of PKD2L1+ cells per section differed significantly among suppliers (F(2, 24) = 32.74, p < 0.0001; Fig. 2D). Jackson mice had more PKD2L1+ cells than Charles River (p = 0.0002) and Taconic mice (p < 0.0001), while Charles River mice had more cells than Taconic mice (p = 0.0089). Cell abundance was also greater in cervical than lumbar sections (F(1, 24) = 7.95, p = 0.0095). Neither the main effect of sex nor the supplier x sex interaction was significant.

The number of PKD2L1+ cells contacting the central canal also differed among suppliers (F(2, 24) = 67.30, p < 0.0001; Fig. 2E). Jackson mice had more contacting cells than Charles River and Taconic mice (both p < 0.0001), whereas Charles River and Taconic mice did not differ significantly (p = 0.065). Cervical sections contained more contacting cells than lumbar sections (F(1, 24) = 7.33, p = 0.012). We also observed a significant supplier x sex interaction (F(2, 24) = 8.60, p = 0.0015). Follow-up comparisons revealed that Jackson males had more canal-contacting cells than Jackson females (Holm-adjusted p = 0.0004), whereas within-supplier sex differences were not significant for Charles River or Taconic mice.

The proportion of PKD2L1+ cells contacting the central canal differed among suppliers (F(2, 24) = 11.26, p = 0.0004; Fig. 2F), with a significant supplier x sex interaction (F(2, 24) = 7.92, p = 0.0023). Averaged across sexes, Jackson mice had a greater contacting proportion than Charles River (p = 0.0015) and Taconic mice (p = 0.0009). Sex-stratified comparisons revealed significant supplier differences among males but not females. Within Taconic mice, females had a greater contacting proportion than males (Holm-adjusted p = 0.011). There was not a significant main effect of spinal region (p = 0.58).

Finally, average soma distance from the central canal differed significantly among suppliers (F(2, 24) = 4.54, p = 0.021; Fig. 2G), with a significant supplier x sex interaction (F(2, 24) = 6.95, p = 0.0042). Although the sex-marginal comparison indicated shorter soma distances in Jackson than Taconic mice (Tukey-adjusted p = 0.024), sex-stratified comparisons revealed that this difference was significant among males (p = 0.0003) but not females (p = 0.83). Within Taconic mice, males had greater soma distances than females (Holm-adjusted p = 0.016). There was not a significant main effect of spinal region (p = 0.068).

These findings demonstrate supplier-associated differences in CSF-cN abundance and organization relative to the central canal, with additional sex-dependent variation in canal contact and soma position.

### CSF-cN abundance and central canal contact differ between cervical and lumbar spinal cord

We next examined regional differences in CSF-cN abundance and organization by directly comparing cervical and lumbar measurements within the same animals (Fig. 3). The cervical spinal cord contained significantly more PKD2L1+ cells per section than the lumbar spinal cord (Fig. 3A; Wilcoxon signed-rank test, W = 98, p = 0.0047). The number of PKD2L1+ cells contacting the central canal was also greater in the cervical region (Fig. 2B; W = 88, p = 0.0022).

The proportion of PKD2L1+ cells contacting the central canal showed a trend toward being greater in the cervical than lumbar spinal cord (Fig. 3C; W = 143, p = 0.067), but was not statistically significant. The distance of PKD2L1+ cell somata from the central canal did not differ significantly between regions (Fig. 3D; W = 164, p = 0.16). Thus, the most pronounced regional differences were in CSF-cN abundance and the absolute number of cells contacting the central canal.

**Figure 3.**
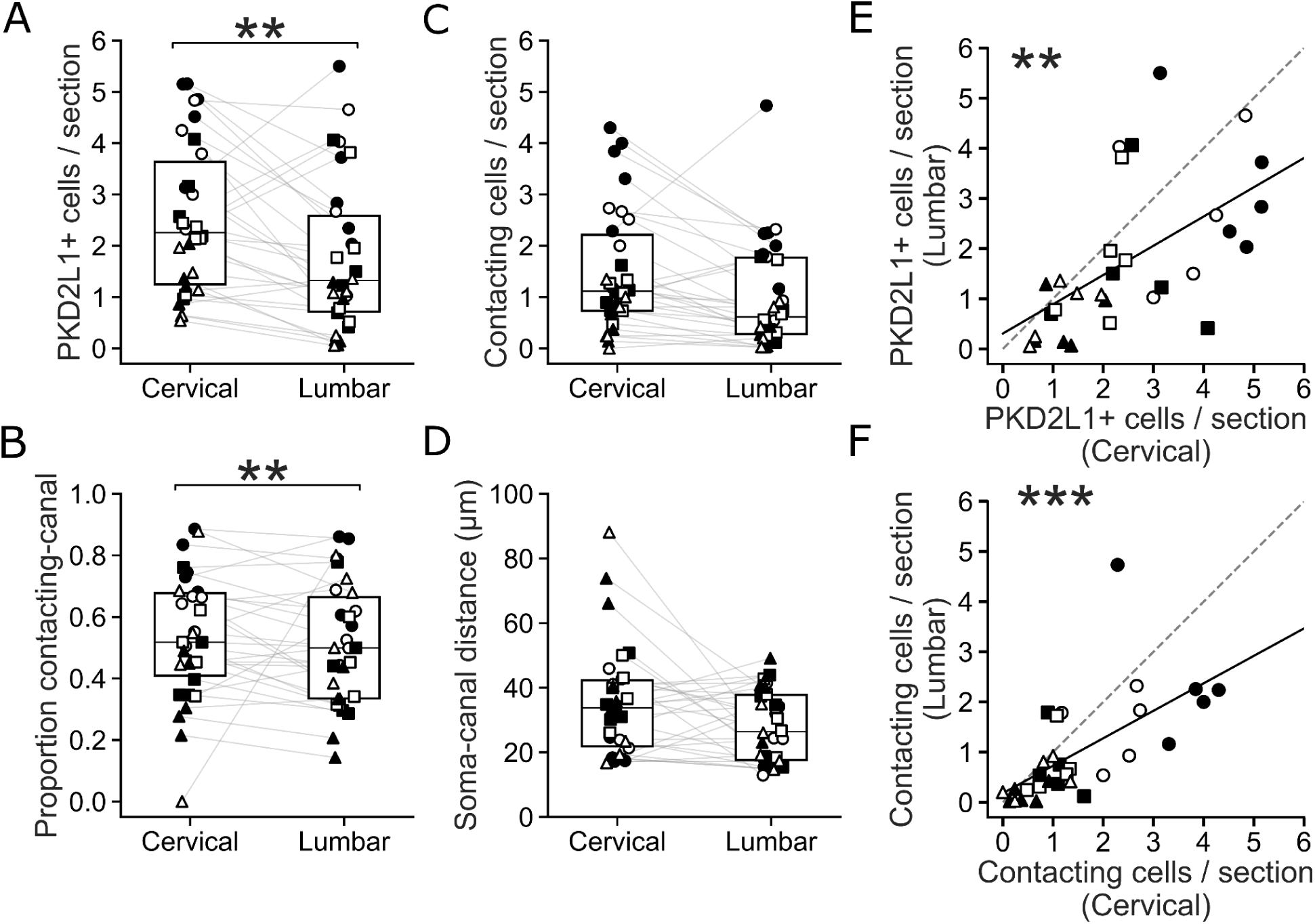
Regional differences in CSF-cN abundance and organization between cervical and lumbar spinal cord. (A-D) Cervical and lumbar measurements were compared within the same animals. (A) Average number of PKD2L1+ cells per section. (B) Average number of PKD2L1+ cells per section contacting the central canal. (C) Proportion of PKD2L1+ cells contacting the central canal. (D) Average distance of PKD2L1+ cell somata from the central canal. (E-F) Relationships between cervical and lumbar measurements across individual animals. (E) Cervical versus lumbar PKD2L1+ cell number. (F) Cervical versus lumbar number of PKD2L1+ cells contacting the central canal. Box plots show the median (Q2), first quartile (Q1), and third quartile (Q3), with individual animals shown as overlaid symbols. Gray lines connect cervical and lumbar measurements from the same mouse in panels A-D. Jackson mice are represented by circles, Charles River mice by squares, and Taconic mice by triangles; male mice are shown with filled symbols and female mice with open symbols. The dashed line has a slope of 1.0. The solid line represents the best fit to the data points. Asterisks indicate statistical significance: *p < 0.05, **p < 0.01, and ***p < 0.001.

CSF-cN measurements were also positively correlated between spinal regions across individual animals (Fig. 3E-F). Mice with greater PKD2L1+ cell abundance in the cervical spinal cord tended to also have greater abundance in the lumbar spinal cord (Fig. 3E; linear regression, slope = 0.58, R² = 0.33, p = 0.0010). Similarly, the number of central canal-contacting PKD2L1+ cells was positively correlated between cervical and lumbar spinal cord (Fig. 3F; slope = 0.55, R² = 0.40, p = 0.0002). These relationships indicate that, despite the overall regional difference in abundance, relative differences among individual animals are shared across cervical and lumbar spinal levels.

### Associations between CSF-cN anatomical measures and motor performance

Finally, we tested whether individual variation in CSF-cN anatomy was associated with performance on the beam, grid, and rotarod tasks (Fig. 4A-I; Table 1). Across the nine comparisons, the relationships were generally weak, with all R² values below 0.10. Total PKD2L1+ cell number was nominally associated with balance beam performance (R² = 0.096, p = 0.016) and rotarod performance (R² = 0.078, p = 0.03), whereas its relationship with grid performance was not significant (R² = 0.044, p = 0.11). Neither the number of PKD2L1+ cells contacting the central canal nor the average distance of cells from the canal was significantly associated with performance on any of the three behavioral tasks. After applying a Bonferroni correction to account for multiple comparisons, none of the nine relationships remained statistically significant. Thus, although suppliers differed in both CSF-cN anatomy and motor performance, individual variation in these anatomical measures showed little correspondence with behavioral performance.

**Figure 4.**
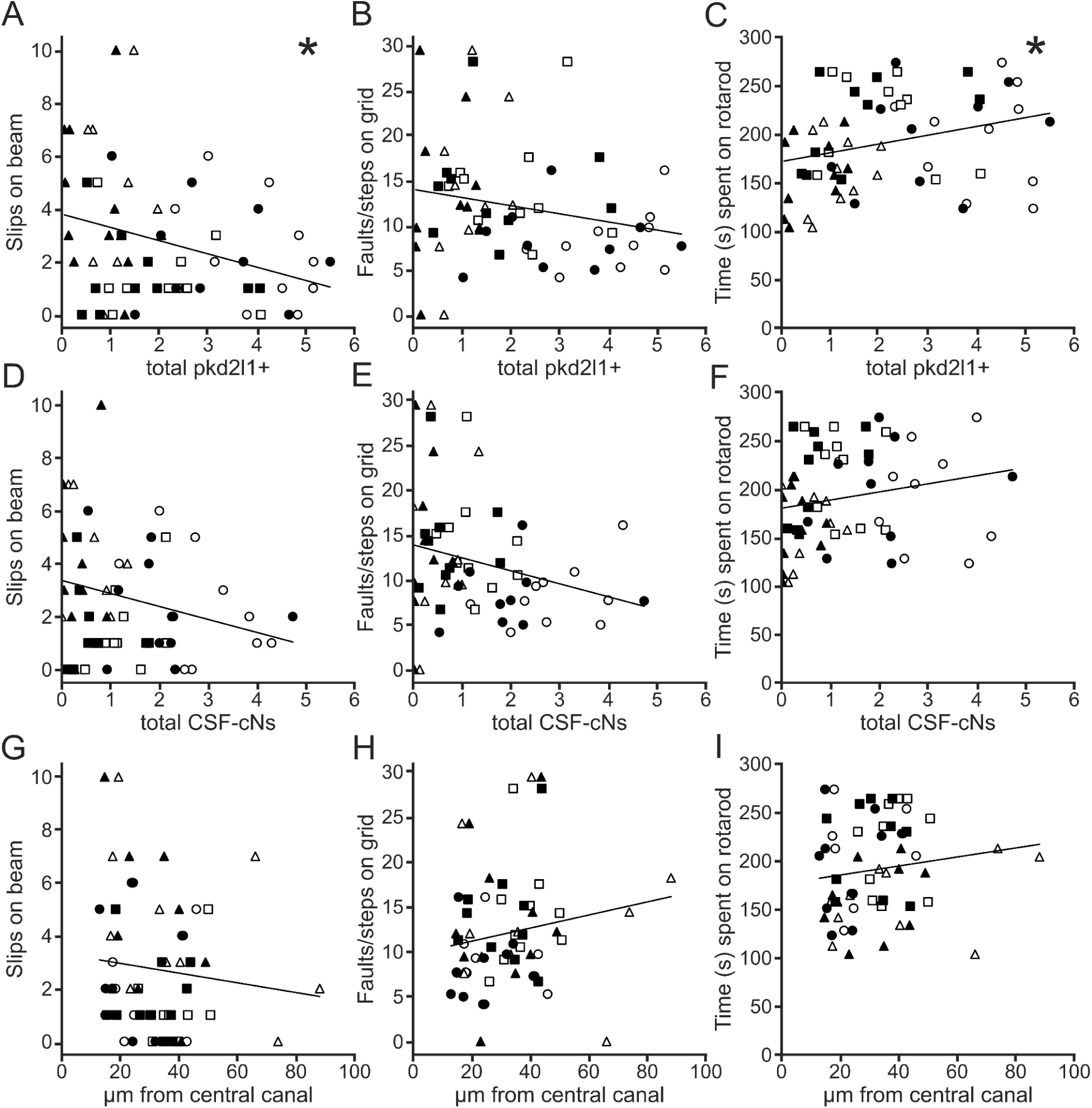
Regression analysis comparing the number of PKD2L1+ cells, the number of cells contacting the central canal and the distance of cells from the central canal with beam, grid, and rotarod performance. (A-C) Average number of PKD2L1+ cells/section plotted against the respective behavioral assessments. (D-F) Average number of PKD2L1+ cells contacting the central canal per section plotted against the respective behavioral assessments. (G-I) Distance of pkd2l1 cell somata from the central canal plotted against the respective behavioral assessments. Jackson mice are represented with circles, Charles River mice by squares, and Taconic mice by triangles. Cervical data are represented by open symbols and lumbar data by closed symbols. Numerical results of the regression analysis are presented in Table 1. Asterisks indicate significant pairwise differences following Tukey’s HSD test *p < 0.05.

**Table 1.**
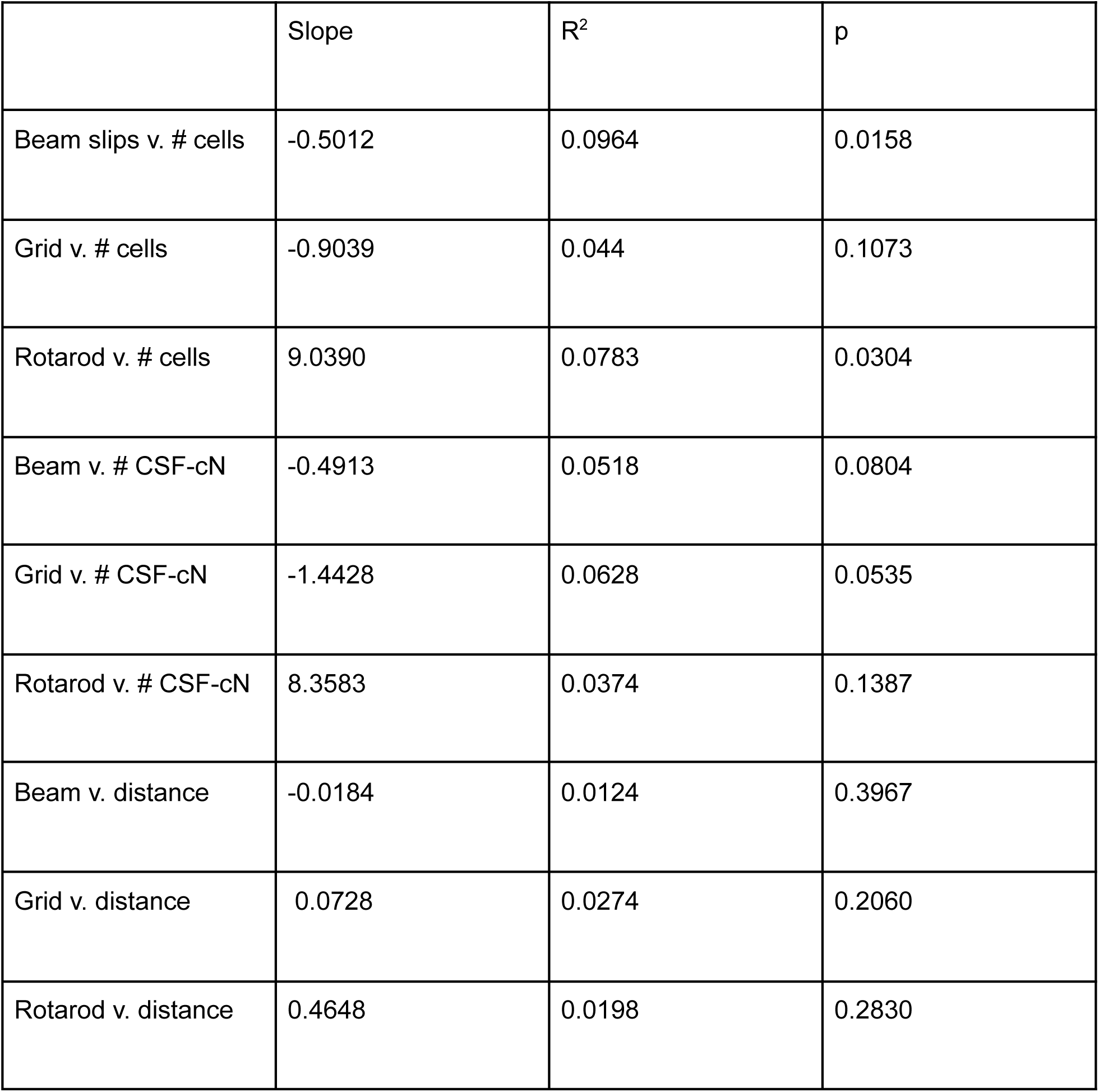
Numerical results for the regression analysis performed on the data presented in Fig. 4.

## DISCUSSION

We found that motor performance and spinal CSF-cN organization vary among closely related C57BL/6 mice obtained from different commercial suppliers. Charles River mice performed better than Taconic mice on the balance beam and rotarod tasks, whereas grid walking performance did not differ significantly among suppliers (Fig. 1A-C). CSF-cN organization also differed substantially among these populations. Jackson mice had the greatest abundance of PKD2L1+ cells, Charles River mice were intermediate, and Taconic mice generally had the fewest. Jackson mice also had more canal-contacting cells and a greater proportion of PKD2L1+ cells contacting the central canal (Fig. 2D-G). Across spinal regions, cervical sections contained more PKD2L1+ cells and more canal-contacting cells than lumbar sections, while the proportion contacting the canal showed only a trend toward a regional difference and soma distance did not differ (Fig.3A-F). Despite supplier-associated differences in both motor performance and CSF-cN organization, individual CSF-cN measures did not show any significant relationships with behavioral performance after applying Bonferroni Correction (Fig. 4).

Consistent with previous reports of behavioral variation among C57BL/6 populations, we observed supplier-associated differences in motor performance. Charles River mice performed better than Taconic mice on the balance beam and rotarod tasks, whereas grid performance did not differ significantly among suppliers. Bryant et al. (2008) also reported differences in rotarod performance, with C57BL/6J males exhibiting longer latencies to fall than C57BL/6NCrl and C57BL/6NTac males, and C57BL/6NCrl males outperforming C57BL/6NTac males. However, the rotarod protocols differed substantially between studies. Bryant et al. accelerated the rotarod from 5 to 60 rpm over 3 min (0.31 rpm/s), whereas our protocol accelerated from 4 to 40 rpm over 5 min (0.12 rpm/s). Bryant et al. also conducted two trials separated by 30 min and repeated testing one week later, using exclusively male mice, whereas our study included both males and females. Differences in acceleration rate, task demands, and testing schedules may influence the relative performance of C57BL/6 populations, although their contributions to the observed differences cannot be determined from these studies.

The supplier-associated anatomical differences suggest that variation among C57BL/6 populations involves more than a simple increase or decrease in CSF-cN abundance (Fig. 2). Jackson mice not only had more PKD2L1+ cells, but also a greater proportion of those cells contacted the central canal (Fig. 3). If supplier-associated variation were restricted to the overall abundance of this population, the relative organization of cells around the canal could remain similar despite differences in cell number. Instead, our results suggest that both the abundance and central canal organization of CSF-cNs vary among these populations. Previous work provides evidence that CSF-cN positioning can differ markedly among C57BL/6 mice. Tonelli Gombalová et al. (2020) reported that the majority of PKD2L1+ CSF-cNs examined in C57BL/6N mice occupied an ectopic ventral position, whereas ectopic cells represented only a small population in C57BL/6J mice. The phenotype described by Tonelli Gombalová et al. (2020) is distinct from the variation measured here, but both studies indicate that CSF-cN anatomy is not uniform across closely related C57BL/6 populations. An important question raised by these findings is which developmental or cellular processes determine both the number of CSF-cNs and their positioning relative to the central canal.

The origins of these supplier-associated differences remain unclear. C57BL/6 populations maintained by different suppliers have distinct breeding histories and can differ genetically, physiologically, and behaviorally (Bryant et al., 2008; Mekada et al., 2009; Mekada and Yoshiki, 2021). Genetic and potentially epigenetic variation, as well as differences in environmental exposures, could contribute to the observed phenotypes. One approach to investigating these differences would be to determine when the anatomical variation first emerges. Comparing CSF-cN anatomy across developmental stages could establish whether supplier-associated differences are present during the formation of the spinal CSF-cN population or develop later. Furthermore, the close genetic relationship among these C57BL/6 populations may provide a useful system for identifying cellular and molecular mechanisms underlying both CSF-cN abundance and organization and variation in gait and balance. Comparative genomic and epigenomic analyses could help identify candidate pathways involved in these phenotypes, while subsequent experimental studies could determine whether the anatomical and behavioral differences arise through shared or independent mechanisms. Our findings do not establish the genetic, epigenetic, or environmental contributions to the observed variation, but highlight how future studies might approach such questions.

Previous studies have not reported a consistent CSF-cN organization along the spinal cord. Liu et al. (2023) found no significant difference between cervical and lumbar spinal cord in the number of PKD2L1+ dendritic bulbs protruding into the central canal, whereas Crozat et al. (2025) reported a greater density of CSF-cNs in lumbar than cervical spinal cord. In contrast, we found greater PKD2L1+ cell abundance in the cervical than lumbar spinal cord, along with a greater number of canal-contacting cells and a trend toward a greater proportion of cells contacting the canal. Although the current anatomical studies do not reveal a consistent spinal region relationship, there is additional evidence that regional CSF-cNs differ functionally. Crozat et al. (2025) found that cervical and lumbar CSF-cNs predominantly exhibited tonic firing, whereas thoracic CSF-cNs more frequently displayed single-spike responses. We did not examine CSF-cN anatomy in the thoracic region, but future studies incorporating this region could provide a more comprehensive understanding of anatomical variation along the spinal cord. These observations highlight that CSF-cNs should not be treated as an anatomically and functionally homogeneous population between spinal regions. Directly examining anatomy and physiology across multiple spinal levels within the same animals could determine whether these forms of regional variation are related.

Although CSF-cN anatomical organization and motor performance both varied among suppliers, the anatomical measurements were not consistently associated with behavioral performance across individual mice. All nine anatomy-behavior regressions accounted for less than 10% of the variance in performance. Total PKD2L1+ cell abundance was associated with beam and rotarod performance before correction for multiple comparisons, but these relationships did not remain significant after correction. The number of canal-contacting cells and soma distance were also not significantly associated with performance. These results address a different question from previous perturbation studies. Ablation or chemogenetic inhibition tests whether CSF-cN activity contributes to locomotor function, whereas our analyses tested whether naturally occurring variation in CSF-cN anatomy covaries with performance among intact animals. Thus, the established functional role of CSF-cNs does not necessarily predict a relationship between cell quantity and motor performance (Gerstmann et al., 2022; Nakamura et al., 2023). One possibility is that the relationship between CSF-cN abundance and locomotor function is nonlinear. Previous work suggests that reductions in CSF-cN number or function do not affect all aspects of locomotion equally. For example, studies involving ablation or inhibition of CSF-cNs impairs skilled locomotion while leaving general locomotor activity and basic gait largely intact, and compensatory sensorimotor pathways have been proposed to maintain locomotion when CSF-cN function is reduced (Gerstmann et al., 2022; Nakamura et al., 2023). These findings raise the possibility that motor performance may not scale proportionally with CSF-cN abundance. If a sufficient population of CSF-cNs can provide the sensory feedback required for normal circuit function, additional neurons may provide little further behavioral advantage. Under this hypothesis, the intermediate abundance observed in Charles River mice could be sufficient to support effective CSF-cN function, while other physiological or circuit properties could contribute more strongly to variation in performance. Our data do not establish a threshold or saturating relationship, but experimental manipulation of the number or recruitment of functional CSF-cNs could directly test this possibility.

Overall, our findings demonstrate that commercially sourced C57BL/6 mice differed in both motor performance and the organization of the CSF-cN population. Supplier-associated variation involved CSF-cN abundance and central canal organization, while cervical and lumbar spinal cord differed most clearly in CSF-cN abundance. Greater CSF-cN abundance did not correspond directly to better motor performance. Instead, physiological and circuit-level properties of CSF-cNs, nonlinear relationships between neuronal abundance and function, or variation elsewhere in the sensorimotor system may contribute to the behavioral differences among these populations. These findings emphasize commercial source as an important consideration in studies of spinal sensorimotor circuitry and provide a framework for investigating the mechanisms that generate variation in the CSF-cN population.

## MATERIALS AND METHODS

### Animals

Adult male and female C57BL/6 mice were obtained at 11 weeks of age from The Jackson Laboratory (Jackson), Charles River Laboratories (Charles River), and Taconic Biosciences (Taconic). A total of 30 mice were used, with 10 mice from each supplier (5 males and 5 females). Mice were housed in same-sex groups according to the commercial supplier, with five animals per cage. Food and water were available ad libitum. Animals were maintained under the animal facility’s standard light-dark cycle. Behavioral testing was conducted over two weeks when mice were 17-18 weeks old. All animals were tested each day in randomized order. All procedures were approved by the University of Washington Institutional Animal Care and Use Committee.

### Behavioral Testing

Behavioral testing was carried out in the Behavioral Core Facility of the Institute for Human Development and Disability.

### Balance Beam

Mice were trained for 5-6 trials on a 25-mm-wide beam. Test trials were performed on a 12-mm-wide aluminum beam. The number of foot faults was counted over a 40-cm section of the beam, excluding the starting area and the portion immediately before the goal box.

### Wire Grid

Untrained mice were placed on a 14 x 12 wire grid with 1-in square openings for 5 min. The total number of hindlimb steps and the number of hindlimb foot faults were counted. Grid performance was calculated as the percentage of hindlimb steps that resulted in a foot fault.

### Rotarod

Mice were trained on a rotarod (ENV-577M; Med Associates Inc., Georgia, VT) at a fixed speed of 16 rpm for 5 min. During test trials, mice were placed on the rod at an initial speed of 4 rpm, which increased to 40 rpm over 5 min. The time each mouse remained on the rod was recorded.

### Histology

#### Tissue Preparation

We anesthetized mice with isoflurane and euthanized them by transcardial perfusion with saline followed by 4% paraformaldehyde in 0.1M phosphate buffer (PB). The spinal column was post-fixed in 4% paraformaldehyde in phosphate buffer (PB) for 24 h at 4C. We dissected the entire spinal cord from the bone using forceps and scissors, then placed it in 20% sucrose in PB for at least overnight. Then we transversely sectioned both the cervical and lumbar regions of the spinal cord at 30 µm thickness on a cryostat (Thermo Scientific HM525 NX) in alternating series with control slides.

### Immunohistochemistry

We dried sections for 30 minutes at room temperature (RT), then washed 3x in 0.1% Triton X-100 in PBS (PBST) before applying a 5% normal goat serum (Abcam or Vector Labs) as a blocker. We incubated sections with primary antibody (Ab) overnight at 4C. We then washed the sections 3x in PBST, incubated them with secondary Ab for 1.5 h - 2 h at RT, and washed 2x in PBST, PBS and dH_2_O. We incubated sections in aqueous DAPI (Biotum; 1:1000) for 10 minutes before coverslipping with Fluoromount G (SouthernBiotech).

Primary Abs were rabbit anti-Polycystin-L (Pkd2L1; EMD Millipore AB9084, Lot # 4019801) diluted 1:500 in PBS. Secondary Abs were AlexaFluor goat-anti rabbit 647 and goat-anti rabbit 488 (Invitrogen; 1:500).

### Cell Quantification

We manually counted PKD2L1+ cells using an epifluorescence microscope (Nikon Eclipse E6000). A PKD2L1+ cell was classified as contacting the central canal when either the cell body or a cellular process visibly contacted the canal. The proportion of PKD2L1+ cells contacting the central canal was calculated by dividing the number of contacting cells by the total number of PKD2L1+ cells.

The distance of each PKD2L1+ cell from the central canal was measured from the center of the cell body to the inner wall of the central canal using ImageJ2 (2.14.0/1.54f). Images were calibrated at 6.14 pixels/µm. We averaged measurements across sections within each spinal region for each mouse before statistical analysis.

### Statistical Analysis and Software

Statistical significance was defined as p < 0.05 unless otherwise specified. Statistical analyses were performed using custom Python scripts (Python 3.11.5). Analyses were performed using pandas (2.2.2), NumPy (1.26.4), SciPy (1.13.1), and statsmodels (0.14.2). Figures were generated using Matplotlib (3.9.2), seaborn (0.13.2), and Inkscape (1.4.3).

To compare motor performance among commercial suppliers, we performed separate one-way analyses of variance (ANOVAs) for balance beam, grid, and rotarod performance, followed by Tukey’s honestly significant difference (HSD) tests for pairwise comparisons (Fig. 1). We additionally examined the effects of supplier and sex on behavioral performance using two-way ANOVAs.

To examine differences in CSF-cN anatomy, we performed repeated-measures ANOVAs for four anatomical variables: the number of PKD2L1+ cells per section, the number and proportion of cells contacting the central canal, and average soma distance from the central canal (Fig. 2). Supplier and sex were treated as between-subject factors, and spinal region (cervical or lumbar) was treated as a within-subject factor. Each mouse served as the unit of repeated measurement. We measured two spinal regions per mouse, so between-subject effects were evaluated using animal-level means across regions, while spinal-region effects and their interactions were evaluated using within-animal differences. Factorial ANOVAs used Type III sums of squares with sum-to-zero contrasts. Supplier comparisons were evaluated using model-based Tukey-adjusted pairwise tests. For significant supplier x sex interactions, we performed sex-stratified supplier comparisons using Tukey adjustment within each sex and within-supplier sex comparisons using Holm adjustment across suppliers.

To compare cervical and lumbar anatomical measurements within individual mice, we used two-sided Wilcoxon signed-rank tests, with Holm correction for the four anatomical comparisons (Fig. 3). We performed ordinary least-squares linear regression to examine relationships between cervical and lumbar measurements of total PKD2L1+ cell abundance and canal-contacting cell abundance.

We used simple linear regression to examine associations between CSF-cN anatomical measurements and behavioral performance (Fig. 4). Total PKD2L1+ cell abundance, the number of canal-contacting cells, and average soma distance were each evaluated against performance on the balance beam, grid, and rotarod tasks, resulting in nine comparisons. A Bonferroni correction was applied across these comparisons.

## REFERENCES

Agduhr E (1922) Über ein zentrales Sinnesorgan (?) bei den Vertebraten. Zeitschrift für Anatomie und Entwicklungsgeschichte 66:223–360.

Böhm UL, Prendergast A, Djenoune L, Nunes Figueiredo S, Gomez J, Stokes C, Kaiser S, Suster M, Kawakami K, Charpentier M, Concordet J-P, Rio J-P, Del Bene F, Wyart C (2016) CSF-contacting neurons regulate locomotion by relaying mechanical stimuli to spinal circuits. Nature Communications 7:10866.

Bryant CD, Zhang NN, Sokoloff G, Fanselow MS, Ennes HS, Palmer AA, McRoberts JA (2008) Behavioral Differences among C57BL/6 Substrains: Implications for Transgenic and Knockout Studies. Journal of Neurogenetics 22:315–331.

Crozat E, Blasco E, Ramirez-Franco J, Riondel P, Jurčić N, Seddik R, Michelle C, Trouslard J, Wanaverbecq N (2025) Cerebrospinal fluid-contacting neurons are sensory neurons with uniform morphological and region-specific electrophysiological properties in the mouse spinal cord. Communications Biology 8:1233.

Delmas P (2004) Polycystins: From Mechanosensation to Gene Regulation. Cell 118:145–148.

Djenoune L, Khabou H, Joubert F, Quan FB, Nunes Figueiredo S, Bodineau L, Del Bene F, Burcklé C, Tostivint H, Wyart C (2014) Investigation of spinal cerebrospinal fluid-contacting neurons expressing PKD2L1: evidence for a conserved system from fish to primates. Front Neuroanat 8:26.

Djenoune L, Wyart C (2017) Light on a sensory interface linking the cerebrospinal fluid to motor circuits in vertebrates. Journal of Neurogenetics 31:113–127.

Fidelin K, Djenoune L, Stokes C, Prendergast A, Gomez J, Baradel A, Del Bene F, Wyart C (2015) State-Dependent Modulation of Locomotion by GABAergic Spinal Sensory Neurons. Curr Biol 25:3035–3047.

Gerstmann K, Jurčić N, Blasco E, Kunz S, de Almeida Sassi F, Wanaverbecq N, Zampieri N (2022) The role of intraspinal sensory neurons in the control of quadrupedal locomotion. Current Biology 32:2442–2453.e4.

Hubbard JM, Böhm UL, Prendergast A, Tseng P-EB, Newman M, Stokes C, Wyart C (2016) Intraspinal Sensory Neurons Provide Powerful Inhibition to Motor Circuits Ensuring Postural Control during Locomotion. Current Biology 26:2841–2853.

Jalalvand E, Robertson B, Wallén P, Grillner S (2016) Ciliated neurons lining the central canal sense both fluid movement and pH through ASIC3. Nat Commun 7:10002.

Kolmer W (1931) Über das Sagittalorgan, ein zentrales Sinnesorgan der Wirbeltiere, insbesondere beim Affen. Zeitschrift für Zellforschung und Mikroskopische Anatomie 13:236–248.

Liu X, Rich K, Nasseri SM, Li G, Hjæresen S, Finsen B, Scherberger H, Svenningsen Å, Zhang M (2023) A Comparison of PKD2L1-Expressing Cerebrospinal Fluid Contacting Neurons in Spinal Cords of Rodents, Carnivores, and Primates. IJMS 24:13582.

Mekada K, Abe K, Murakami A, Nakamura A, Nakata H, Moriwaki K, Obata Y, Yoshiki A (2009) Genetic Differences among C57BL/6 Substrains. Experimental Animals 58:141–149.

Mekada K, Yoshiki A (2021) Substrains matter in phenotyping of C57BL/6 mice. Experimental Animals 70:145–160.

Nakamura Y, Kurabe M, Matsumoto M, Sato T, Miyashita S, Hoshina K, Kamiya Y, Tainaka K, Matsuzawa H, Ohno N, Ueno M (2023) Cerebrospinal fluid-contacting neuron tracing reveals structural and functional connectivity for locomotion in the mouse spinal cord. Elife 12.

Orts-Del’Immagine A, Wanaverbecq N, Tardivel C, Tillement V, Dallaporta M, Trouslard J (2012) Properties of subependymal cerebrospinal fluid contacting neurones in the dorsal vagal complex of the mouse brainstem. The Journal of Physiology 590:3719–3741.

Tonelli Gombalová Z, Košuth J, Alexovič Matiašová A, Zrubáková J, Žežula I, Giallongo T, Di Giulio AM, Carelli S, Tomašková L, Daxnerová Z, Ševc J (2020) Majority of cerebrospinal fluid-contacting neurons in the spinal cord of C57Bl/6N mice is present in ectopic position unlike in other studied experimental mice strains and mammalian species. Journal of Comparative Neurology 528:2523–2550.

Vitar M, Prieto D, Malas S, Russo RE, Trigo FF (2026) PKD2L1 channels segregated to the apical compartment are the dual-mode pH sensor in cerebrospinal fluid-contacting neurons. eLife 15:RP109372.

Wu M-Y, Carbo-Tano M, Mirat O, Lejeune F-X, Roussel J, Quan FB, Fidelin K, Wyart C (2021) Spinal sensory neurons project onto the hindbrain to stabilize posture and enhance locomotor speed. Current Biology 31:3315–3329.e5.

Wyart C, Carbo-Tano M, Cantaut-Belarif Y, Orts-Del’Immagine A, Böhm UL (2023) Cerebrospinal fluid-contacting neurons: multimodal cells with diverse roles in the CNS. Nature Reviews Neuroscience 24:540–556.

